# Jasmonate-induced expansion of the MYC2 cistrome occurs within a preconfigured chromatin landscape

**DOI:** 10.64898/2026.10.01.755910

**Authors:** Linkan Dash, Moonia Ammari, Aanchal Choudhary, Kashif Maseh, Moorthy Gnanarajah, Jeevika Gupta, Mark Zander

## Abstract

The transcription factor (TF) MYC2 is a central regulator of jasmonate (JA) signaling, yet its genome-wide occupancy has primarily been characterized using transgenic lines expressing tagged MYC2 proteins, leaving the dynamics of the native MYC2 cistrome unresolved. Here, using a MYC2-specific antibody, we define the wildtype MYC2 cistrome in *Arabidopsis thaliana* under basal and JA-induced conditions. Native MYC2 showed minimal genome-wide occupancy under control conditions but underwent a pronounced expansion following JA treatment, resulting in more than 3,000 binding sites enriched at core components of the JA signaling network. Cistrome expansion coincided with increased MYC2 protein abundance, while elevated basal MYC2 levels in a transgenic line were associated with extensive MYC2 occupancy even in the absence of JA, supporting a dosage-dependent relationship between MYC2 abundance and genome-wide binding. Despite this extensive cistrome expansion and transcriptional reprogramming, ATAC-seq revealed remarkably limited remodeling of chromatin accessibility following JA treatment. MYC2 recruitment occurred predominantly within pre-existing accessible chromatin regions that remained largely intact in *myc2 myc3 myc4* mutants. Together, our findings reveal that JA-dependent expansion of the native MYC2 cistrome occurs within a largely pre-established accessome and identify MYC2 dosage as an important determinant of the extent of genome-wide occupancy.

## INTRODUCTION

Plants continuously encounter fluctuating environmental conditions and must rapidly adjust gene expression to optimize growth, development, and defense. Central to these adaptive responses are transcription factors (TFs), which integrate environmental signals with chromatin-based regulatory mechanisms to coordinate the establishment and maintenance of gene regulatory networks (Zander and Vesper, 2026). Prime examples of TFs whose activities are governed by external cues are the basic helix-loop-helix (bHLH) TFs MYC2 (MYELOCYTOMATOSIS 2), MYC3, MYC4, and MYC5 (MYCs), which control the jasmonic acid (JA) pathway thereby coordinating diverse aspects of plant growth, development, and responses to biotic and abiotic stresses (Lorenzo et al., 2004, Fernandez-Calvo et al., 2011, Dombrecht et al., 2007, Qi et al., 2015, Schweizer et al., 2013). Current models suggest that, under non-inducing conditions, MYCs are repressed by JAZ (JASMONATE ZIM-DOMAIN) proteins, which connect MYCs with a repressive TOPLESS (TPL) complex comprising NINJA, TPL-related proteins (TPRs), histone deacetylases (HDAs), and GCN5 (GENERAL CONTROL NON-DEPRESSIBLE5) (Chini et al., 2007, Thines et al., 2007, Pauwels et al., 2010, Li et al., 2020, An et al., 2022). Accumulation of the bioactive hormone (+)-7-iso-jasmonoyl-L-isoleucine (JA-Ile) promotes the association of the Skp1-Cullin F-box E3 ubiquitin ligase complex SCF^COI1^ with JAZs to form the JA-Ile co-receptor complex (Thines et al., 2007, Chini et al., 2007, Sheard et al., 2010, Fonseca et al., 2009). This results in JAZ ubiquitination and proteasomal degradation, thereby releasing MYCs to engage with the permissive MED25-HAC1 (HISTONE ACETYLTRANSFERASE OF THE CBP FAMILY1) - LUH (LEUNIG_HOMOLOG) complex and activate target gene expression (Chini et al., 2007, Thines et al., 2007, Wang et al., 2019, An et al., 2017, You et al., 2019).

This transcriptional activation is accompanied by extensive reprogramming of the chromatin landscape at JA-responsive genes, both across gene bodies and in broad regions surrounding MYC2 binding sites, a chromatin feature that we refer to as SIENA (Stimulus-Induced Enhancer Acetylation) domains (Choudhary et al., 2024, Zander and Vesper, 2026, Wang et al., 2019, An et al., 2017, You et al., 2019). Notably, loss of MYC activity strongly impairs both epigenomic and transcriptional reprogramming and consequently compromises immune responses against herbivorous insects and fungal pathogens (Zander et al., 2020, Choudhary et al., 2024, Lorenzo et al., 2004, Dombrecht et al., 2007, Fernandez-Calvo et al., 2011, Schweizer et al., 2013). The central role of MYCs in plant immunity is further underscored by recent findings showing that increasing MYC2 dosage is critical for conferring citrus Huanglongbing resistance (Zhao et al., 2025). Thus, a comprehensive understanding of the molecular mechanisms underlying MYC function is important for developing strategies to improve stress resilience in crops, beginning with a comprehensive characterization of their repertoire of DNA-binding sites - their cistromes.

The first insights into the MYC2 cistrome were obtained almost 30 years ago (Abe et al., 1997, dePater et al., 1997). *Arabidopsis* MYC2 binds to the canonical G-box (5′-CACGTG-3′) as well as numerous sequence variants, a binding preference that was first identified *in vitro* using electrophoretic mobility shift assays (EMSAs) (Abe et al., 1997, Boter et al., 2004, Yadav et al., 2005, dePater et al., 1997) and subsequently confirmed using CELD-fusion DNA-binding assays and protein-binding microarrays (Dombrecht et al., 2007, Godoy et al., 2011). The first *in vivo* evidence for MYC2 DNA binding was obtained by ChIP-qPCR (chromatin immunoprecipitation coupled with quantitative real-time PCR) demonstrating MYC2 occupancy at the promoters of *LOX2* (*LIPOXYGENASE2*) and *TAT1* (*TYROSINE AMINOTRANSFERASE1*) in *Arabidopsis* (Hou et al., 2010). Subsequently, ChIP-seq (chromatin immunoprecipitation followed by sequencing) enabled genome-wide profiling of MYC2 DNA binding, generating MYC2 cistromes in *Arabidopsis*, tomato, and rice (Wang et al., 2019, Van Moerkercke et al., 2019, Zander et al., 2020, Choudhary et al., 2024, Du et al., 2017, Chen et al., 2024, Schweizer et al., 2013). Moreover, CUT&Tag (cleavage under targets and tagmentation) has been used to generate MYC2 cistromes in maize (Ma et al., 2023).

In addition to *in vivo* cistrome mapping techniques, DAP-seq (DNA affinity purification sequencing) has been widely used to generate *in vitro* cistromes of MYC2 and related bHLH TFs across diverse plant species including ginkgo, poplar, maize, *Marchantia*, tomato, and grapevine (Galli et al., 2025, O’Malley et al., 2016, Liu et al., 2024, Marques-Galvez et al., 2024, Zheng et al., 2024, López-Vidriero et al., 2021, Zhang et al., 2025). Comparative DAP-seq analyses further demonstrate that DNA features beyond the core binding motif contribute to target recognition by MYC2-related proteins (López-Vidriero et al., 2021). These studies have substantially expanded our understanding of the sequence determinants and potential genomic target space of MYC2. Although DAP-seq measures TF-DNA binding *in vitro* and does not capture the effects of endogenous MYC2 abundance or the native chromatin context, it has been instrumental in generating cistrome information for species that are difficult to transform and for which such information would otherwise remain unavailable.

TF cistromes are highly dynamic and shaped by diverse internal and external cues (Ammari et al., 2026a) and especially TFs operating in pathways underlying plant-environment interactions often undergo cue-specific cistrome dynamics that are controlled by changes in their abundance or availability (Willige et al., 2021, Chang et al., 2013, Song et al., 2016). By mapping *in vivo* cistromes, primarily using ChIP-seq in the absence and presence of a stimulus, these cue-specific cistrome dynamics can be captured. This principle is exemplified by EIN3 (Chang et al., 2013), which is stabilized in response to ethylene, resulting in increased genome-wide DNA binding. Similarly, PIF7 is sequestered by phyB in photobodies under white light but dissociates under low red-to-far-red (R:FR) light conditions, allowing PIF7 to bind its genomic targets (Willige et al., 2021). The abundance of WRKY33 also substantially increases following *Botrytis cinerea* infection, establishing a pathogen-induced WRKY33 cistrome (Liu et al., 2015).

A major challenge of these *in vivo* cistrome-mapping approaches is the need for TF-specific antibodies, which remain scarce for plant TFs. The first reported EIN3 ChIP-seq study represents an ideal case, in which an EIN3-specific antibody was used to map endogenous EIN3 occupancy in *Arabidopsis* Col-0 seedlings (Chang et al., 2013), but such examples remain the exception. Instead, most plant TF ChIP-seq studies rely on transgenic lines expressing epitope-tagged TFs. All reported MYC2 *in vivo* cistromes are derived from transgenic, epitope-tagged *MYC2* lines which have potentially an altered MYC2 dosage (Zander et al., 2020, Choudhary et al., 2024, Van Moerkercke et al., 2019, Wang et al., 2019, Du et al., 2017).

A critical mechanistic prerequisite for cistrome expansion is accessible chromatin, as most TFs bind DNA within nucleosome-depleted regions. Although JA signaling rapidly induces extensive transcriptional and epigenomic reprogramming in *Arabidopsis* wildtype plants (Choudhary et al., 2024, Hickman et al., 2017, Zander et al., 2020, Ammari et al., 2026b), the extent to which these responses depend on *de novo* MYC2 DNA binding is not well understood. Moreover, whether regulatory regions of JA-responsive MYC2 target loci are already accessible prior to MYC2 binding has not been systematically investigated. It also remains unclear whether chromatin accessibility facilitates MYC2 binding or whether *MYC2* itself contributes to establishing accessible chromatin.

Here, we captured the wildtype MYC2 cistrome in *Arabidopsis* under basal and JA-induced conditions using a validated native anti-MYC2 antibody. We uncovered a pronounced JA-dependent expansion of MYC2 occupancy. Comparison with transgenic MYC2 cistromes and MYC2 protein abundance reveals a strong relationship between MYC2 dosage and cistrome composition. Surprisingly, ATAC-seq shows that this extensive cistrome expansion occurs with minimal remodeling of chromatin accessibility, with MYC2 recruitment occurring predominantly within pre-existing accessible chromatin regions (ACRs) that are maintained independently of MYC2/3/4 as well as of the *Arabidopsis* BAS (BRAHMA-associated SWI/SNF), SAS (SPLAYED-associated SWI/SNF), and MAS (MINUSCULE1/2-associated SWI/SNF) chromatin remodeling complexes. Together, our findings reveal that JA-induced expansion of the MYC2 cistrome occurs within a largely preconfigured accessible chromatin landscape and support TF abundance as an important determinant of stimulus-dependent genome-wide occupancy

## RESULTS

### Validation of an anti-MYC2 antibody

Previous genome-wide studies have established the *Arabidopsis* MYC2 cistrome using JA-treated transgenic lines expressing tagged MYC2 proteins (Zander et al., 2020, Wang et al., 2019, Choudhary et al., 2024). However, the MYC2 cistrome in Col-0 plants (hereafter referred to as the wildtype cistrome) has not been determined, leaving unresolved whether transgenic approaches accurately capture MYC2 occupancy under non-inducing and high-JA conditions and, importantly, the transition between these states. To establish a wildtype benchmark, we tested the specificity of a commercially available rabbit polyclonal anti-MYC2 antibody and evaluated its suitability for MYC2 cistrome mapping by ChIP-seq. We first performed immunoblot analyses in untreated and JA-treated (2 h) Col-0, *myc2*, and *myc2 myc3 myc4* (*myc234*) axenically grown 10-day-old seedlings. In Col-0, we detected a band at the expected size of MYC2 (∼68 kDa) under control conditions that increased substantially following JA treatment (Fig. 1A, and Supplementary Fig. S1A, B), consistent with the previously reported JA-induced accumulation of MYC2 (Zhai et al., 2013). In contrast, this band was absent in the *myc2* mutant under both conditions (Fig. 1A, and Supplementary Fig. S1A, B). A similar pattern was observed in *myc234* seedlings, although longer exposure revealed a faint band at the expected size of MYC2 (Fig. 1A, and Supplementary Fig. S1A, B). We attribute this signal to the presence of a nonfunctional MYC2 protein encoded by the *jin1-2* allele present in *myc234* (Lorenzo et al., 2004, Fernandez-Calvo et al., 2011). By contrast, the *myc2* single mutant is a T-DNA insertion allele (*jin1-8*) (Lorenzo et al., 2004), and no MYC2 protein was detected (Fig. 1A, and Supplementary Fig. S1A, B). In addition, we performed immunoblotting in three-week-old soil-grown plants and observed substantial accumulation of MYC2 protein following 2 h of JA treatment (Supplementary Fig. S1C, D). We also tested the MYC2 antibody in untreated and JA-treated tomato tissue but did not detect MYC2, suggesting that the antibody does not efficiently recognize tomato MYC2 (Supplementary Fig. S1A, B). To further assess antibody specificity and MYC2 abundance, we examined the previously described transgenic *myc2 pMYC2:MYC2-FLAG* line and a newly generated *myc234 pMYC2:MYC2-4×MYC* line. In both transgenic lines, MYC2 was detected at a slightly higher molecular weight, consistent with the presence of the additional epitope tags (Supplementary Fig. S1A-D). Notably, the difference in MYC2 protein levels between untreated and JA-treated samples was often minimal in both transgenic lines, with untreated samples frequently exhibiting substantially higher MYC2 levels than untreated Col-0 plants (Supplementary Fig. S1A-C). Lastly, our native anti-MYC2-antibody also successfully recognized recombinant MYC2-4xMYC protein (Fig. 1B).

**Figure 1.**
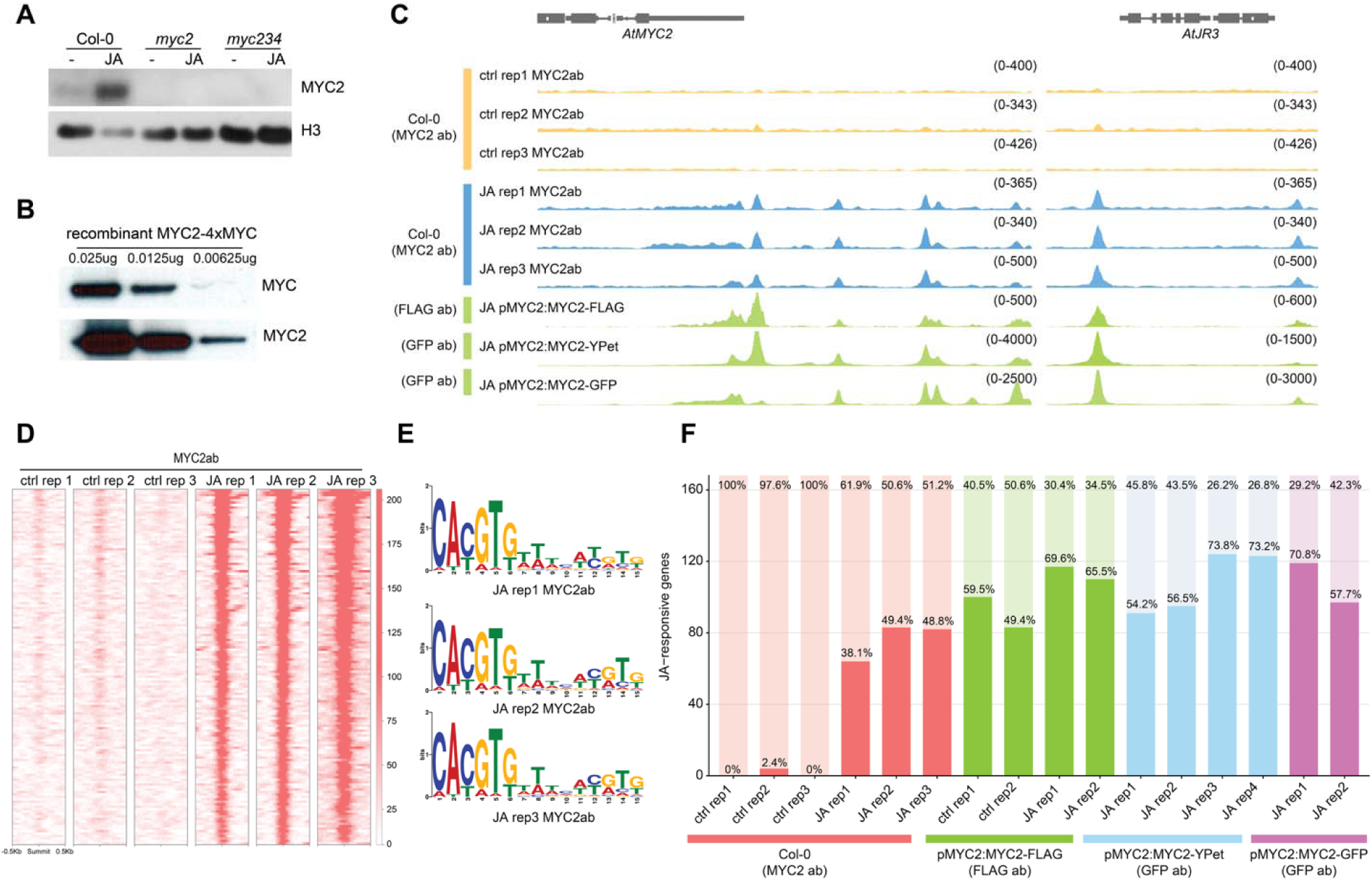
Generation of wildtype MYC2 cistromes. **A**, Immunoblot analysis of the anti-MYC2 antibody using protein extracts from 10-day-old seedlings grown on LS medium. Protein extracts from Col-0, *myc2*, and *myc234*, were analyzed in the absence (−) or presence of JA. Additional replicates are shown as full blots are in Supplementary Figure 1. **B,** Recombinant MYC2-4×MYC protein was analyzed at 0.025, 0,0125, and 0,00625 ug per lane. Protein was detected by immunoblotting using either an anti-MYC antibody (top) or the anti-MYC2 antibody (bottom), confirming detection of recombinant MYC2 by the anti-MYC2 antibody. **C,** Genome browser screenshots of the *MYC2* and *JR3* loci show MYC2 occupancy across indicated experiments. MYC2 occupancy in Col-0 under control conditions and 2 h JA treatment was detected with the native MYC2 antibody (blue). MYC2 occupancy from previously published MYC2 ChIP-seq dataset derived from FLAG-, YPet-, and GFP-tagged MYC2 plants are shown in green. Tracks were normalized to sequencing depth. **D.** Heatmap of summit-centered MYC2 occupancy across 145 target genes in Col-0 control and 2 h JA-treated Col-0 replicates. The 145 gene set was derived by assigning genes to the top 500 MACS2-identified MYC2 peaks and intersecting these peak-associated genes with JA upregulated genes identified by RNA-seq. MYC2 signal is shown within ± 500 bp of peak summits. Heatmap scale color intensity represents MYC2 ChIP-seq signal across the summit centered window. **E**. Representative DNA-binding motifs enriched in native MYC2 ChIP-seq peaks across all here 2 h JA-treated Col-0 replicates. Motif enrichment analysis was performed on the top 500 peak summits selected from MACS2-identified peaks using MEME-ChIP **F.** Bar graph showing the number of genes from a curated set of 168 JA-responsive pathway genes (Zander et al., 2020). Colors indicate the different sets of MYC2 ChIP-seq data used: native MYC2 antibody (red), pMYC2:MYC2-FLAG (green), pMYC2:MYC2-YPet (blue), and pMYC2:MYC2-GFP (purple). Percentages within the bars indicate overlap, while percentages above the bars indicate non-overlap (light-shaded portion).

### Defining the wildtype MYC2 cistrome

Next, we used the anti-MYC2 antibody to map the wildtype MYC2 cistrome in untreated and JA-treated (2 h) Col-0 seedlings using ChIP-seq. In total, we generated three biological replicates for both control and JA-treated samples. Visual inspection of our MYC2 ChIP-seq data in a genome browser revealed clean, distinct peaks across all three JA-treated replicates at the canonical MYC2 target genes *MYC2, JR3, ORA47,* and *JAZ9* (Fig. 1C, and Supplementary Fig. S2A). Similar binding patterns were observed in previously reported MYC2 ChIP-seq datasets generated from *myc2 pMYC2:MYC2-FLAG*, Col-0 *pMYC2:MYC2-YPet*, and Col-0 *pMYC2:MYC2-GFP* plants (Fig. 1C, and Supplementary Fig. S2A) (Zander et al., 2020, Choudhary et al., 2024, Wang et al., 2019). We then identified binding peaks for all replicates and conducted *de novo* motif search of the summits of the top 500 MYC2 peaks (Supplementary Fig. S2B, and Supplementary Table S1). Under control conditions, we identified between 546 and 1,207 peaks; however, these peaks lacked enrichment for the canonical G-box motif, suggesting that they predominantly represent background signal (Supplementary Fig. S2B, and Supplementary Table S1). Thus, no clear peaks were observed at known MYC2 target genes (Fig. 1C). In addition, gene ontology analysis of the control replicates did not reveal any enrichment for JA-regulated processes (Supplementary Fig. S2C). In contrast, 2 h of JA treatment resulted in a pronounced expansion of the MYC2 cistrome, with an average of approximately 3,000 peaks detected genome-wide (Fig. 1C, D, Supplementary Fig. S2B, and Supplementary Table S1). These JA-induced MYC2 peaks showed strong enrichment for the canonical G-box motif, consistent with direct MYC2 binding (Fig. 1E, and Supplementary Fig. S2B).

For comparison, we reanalyzed previously reported MYC2 cistromes generated from JA-treated *myc2 pMYC2:MYC2-FLAG,* Col-0 *pMYC2:MYC2-YPet*, and Col-0 *pMYC2:MYC2-GFP*, transgenic plants (Zander et al., 2020, Choudhary et al., 2024, Wang et al., 2019). The number of detected MYC2 binding sites was substantially higher in all transgenic datasets than in our wildtype cistrome, although considerable variation was also evident among replicates of the published datasets (Supplementary Fig. S2B, and Supplementary Table S1). To assess the comparability of our wildtype cistrome with these previously reported datasets, we next examined MYC2 occupancy across a curated set of 168 core JA pathway genes (Supplementary Table S2) (Zander et al., 2020). This set encompasses major components of the pathway, ranging from JA biosynthesis and metabolism to signaling and transcriptional regulation (Supplementary Table S2) (Zander et al., 2020). Consistent with the extensive feedback regulation characteristic of the JA pathway, MYC2 targeted approximately half of these core components in our wildtype cistrome (Fig. 1F). The transgenic datasets showed a slightly greater number of targeted JA pathway genes ranging from 54% to 73%, but the overall extent of pathway coverage was comparable to that observed with the native MYC2 antibody in Col-0 seedlings (Fig. 1F). Functional annotation of genes comprising the wildtype MYC2 cistrome under JA-inducing conditions further revealed strong enrichment for JA-responsive processes (Supplementary Fig. S2D). Together, these findings support the biological validity of our wildtype MYC2 cistrome and further highlight extensive MYC2-mediated feedback regulation as a central aspect of the JA signaling network.

### MYC2 dosage-dependent expansion of the MYC2 cistrome

A striking feature of our wildtype MYC2 cistrome was its pronounced JA-dependent expansion, which coincided with the strong increase in MYC2 protein abundance following JA exposure (Fig. 1A-D, and Supplementary Fig. S1A-D, S2A, B). Although we detected a small number of peaks under untreated conditions, motif analysis did not identify significant enrichment of the G-box, and none of the core JA pathway components were occupied by MYC2 (Fig. 1F, and Supplementary Fig. S2B). Nevertheless, when averaging the ChIP-seq signal across a set of 145 MYC2 direct target genes which are strongly transcriptionally unregulated by JA (Supplementary Table S1), we detected weak MYC2 occupancy at these sites already under control conditions (Supplementary Fig. S2E), albeit below the peak calling threshold. This is consistent with the low abundance of MYC2 protein in untreated plants (Fig. 1A, B, and Supplementary Fig. S1A-D). Together, the strong JA-induced increase in MYC2 protein abundance and the concomitant expansion of the MYC2 cistrome indicates that MYC2 dosage is a key determinant of genome-wide MYC2 occupancy.

Among the published transgenic datasets, MYC2 cistromes from *myc2 pMYC2-MYC2-FLAG* plants were available under both control and JA conditions, whereas cistromes from Col-0 *pMYC2-MYC2-Ypet* and Col-0 *pMYC2:MYC2-GFP* plants were available only following JA treatment (Wang et al., 2019, Zander et al., 2020, Choudhary et al., 2024). Interestingly, *myc2 pMYC2:MYC2-FLAG* plants already displayed an extensive MYC2 cistrome under control conditions, with 4,057 and 6,306 peaks detected in the two replicates (Supplementary Fig. S2B) Unlike the control cistromes in Col-0, these cistromes were enriched at JA pathway components, and motif analysis of the peak summits revealed strong enrichment of the G-box (Fig. 1F, and Supplementary Fig. S2B), indicating the presence of a bona fide JA-like MYC2 cistrome even in the absence of JA treatment. However, when examining MYC2 occupancy at these sites, we still observed a substantial increase following JA treatment (Supplementary Fig. S2E), indicating that JA strongly enhances MYC2 occupancy even when cistrome composition changes comparatively little. Consistent with this observation, our immunoblot analysis showed substantially higher basal MYC2 protein abundance in Col-0 seedlings (Supplementary Fig. S1A-C). Thus, elevated basal MYC2 levels in *myc2 pMYC2:MYC2-FLAG* plants correlate with extensive MYC2 occupancy already in the absence of JA, further supporting a relationship between MYC2 dosage and cistrome size.

### The *Arabidopsis* accessome is only modestly responsive to JA

The pronounced JA-dependent expansion of the wildtype MYC2 cistrome in Col-0 seedlings prompted us to investigate the extent to which chromatin accessibility contributes to MYC2 cistrome expansion. Apart from pioneer TFs, which remain poorly characterized in plants except LEAFY (Jin et al., 2021), TF binding generally occurs within nucleosome-depleted regions (NDRs). We therefore asked whether JA actively remodels global chromatin accessibility (accessome) to regulate the availability of NDRs for MYC2 binding. To address this question, we performed ATAC-seq (Assay for Transposase-Accessible Chromatin using sequencing) in Col-0 seedlings under control conditions and following 2 h of JA treatment, as well as in JA-treated *myc234* mutant seedlings. In parallel, we profiled the transcriptional response to JA by RNA-seq across a time series comprising control, 30 min, 1 h, 2 h, and 4 h of JA treatment in Col-0 seedlings. As shown previously, the transcriptome is highly responsive to JA (Hickman et al., 2017, Zander et al., 2020), with more than 2000 genes differentially expressed in our time series experiment (Supplementary Fig. S3A-G, and Supplementary Table S3). Typical JA pathway marker genes, including *MYC2* and *JAZ9*, showed robust transcriptional upregulation (Supplementary Fig. S3B, D, E). This rapid and extensive transcriptional response implies that substantial regulatory events, including MYC2 binding, occur prior to or concomitant with gene activation.

Our ATAC-seq analysis yielded 23,569 - 41,004 accessible chromatin regions (ACRs), and integration across biological replicates identified 38,880 high-confidence ACRs (Supplementary Table S4). ATAC-seq fragments overlapping each ACR were quantified and normalized using DESeq2, enabling direct comparison of accessibility between control and JA-treated samples. Differential accessibility analysis identified 1,651 differentially accessible regions (DARs) with significant differences between control and JA-treated Col-0 samples (FDR < 0.05) (Fig. 2A). However, accessibility changes were generally small and predominantly reflected increased accessibility following JA treatment, with none of these DARs exhibiting a ≥2-fold increase (log_₂_FC ≥ 1) (Fig. 2A), indicating that the accessome remains largely stable following JA treatment (Fig. 2A, B). Consistent with these observations, we detected essentially no *de novo* formation of ACRs associated with JA-induced MYC2 recruitment. Of the 1,651 JA-responsive DARs, 244 (14.8%) overlapped MYC2 binding sites, compared with 1,686 of 37,229 (4.5%) non-responsive ACRs, representing an approximately threefold enrichment of MYC2 occupancy at JA-responsive ACRs (Fig. 2C). Thus, although most JA-responsive ACRs were not directly associated with MYC2 binding, MYC2 occupancy was enriched among ACRs exhibiting JA-dependent changes in accessibility (Fig. 2C). In addition, our combined cistrome, transcriptome, and accessome analyses revealed that the pronounced JA-dependent expansion of the MYC2 cistrome and accompanied transcriptome changes occur predominantly within a pre-existing accessome rather than through widespread JA-induced chromatin opening (Fig. 3A).

**Figure 2.**
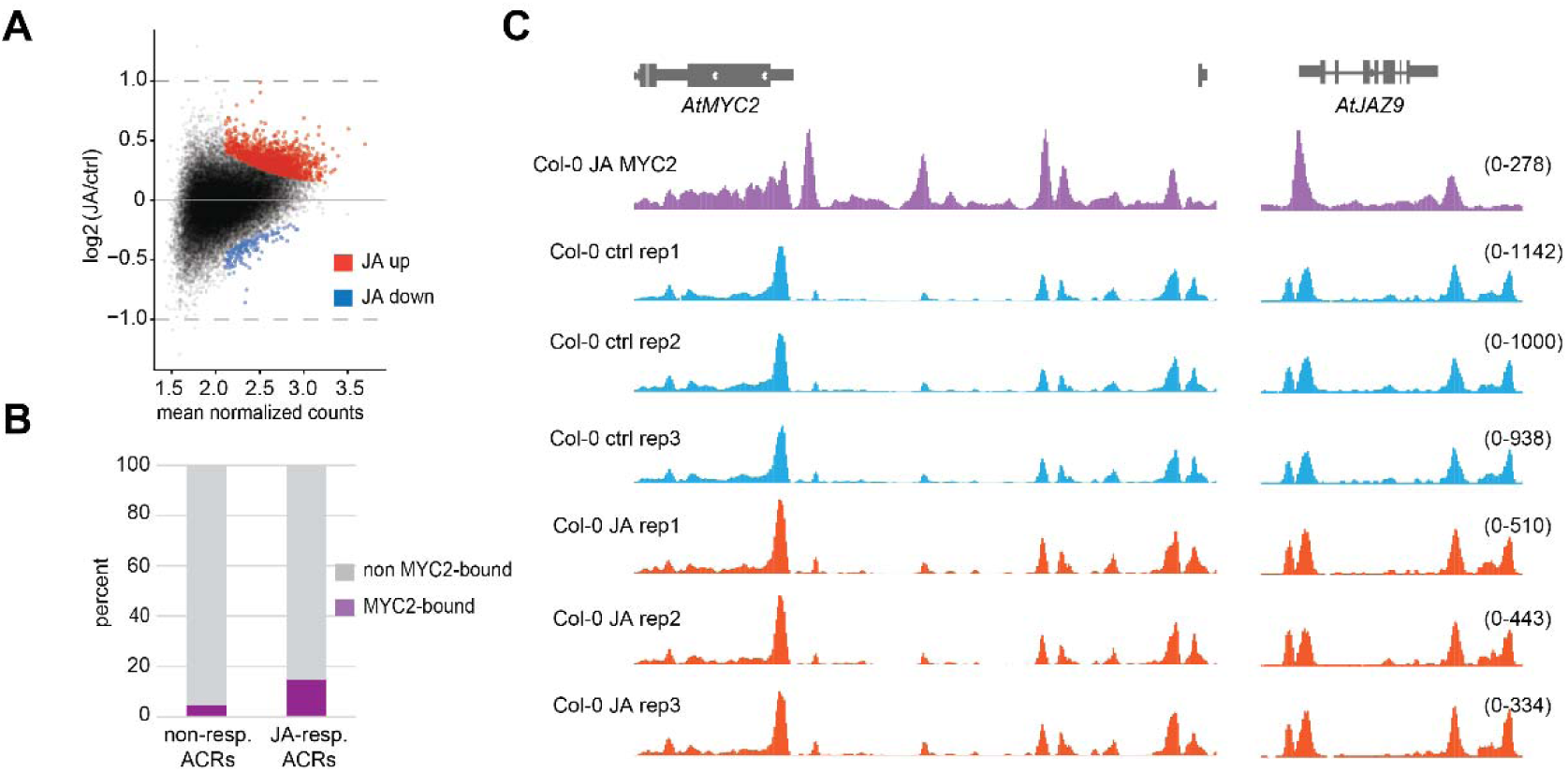
JA only induces very modest changes in chromatin accessibility A, MA plot showing differential chromatin accessibility between control (ctrl) and JA-treated Col-0 seedlings. Each point represents an ACR, plotted according to mean normalized ATAC-seq counts and log2 fold change (JA/ctrl). DARs with significantly increased (red) or decreased (blue) accessibility were identified using DESeq2 (FDR < 0.05). Dashed lines indicate log2 fold changes of ±1. **B,** Percentage of non-responsive and JA-responsive ACRs overlapping MYC2 binding sites. MYC2-bound ACRs are shown in purple and non-MYC2-bound ACRs in gray. **C,** Genome browser screenshots show MYC2 ChIP-seq (JA replicate 1) and ATAC-seq profiles at the *MYC2* and *JAZ9* loci. MYC2 binding following JA treatment is shown in purple. ATAC-seq profiles from three biological replicates of control and JA-treated Col-0 seedlings are shown in blue and orange, respectively. Track scales are indicated on the right.

**Figure 3.**
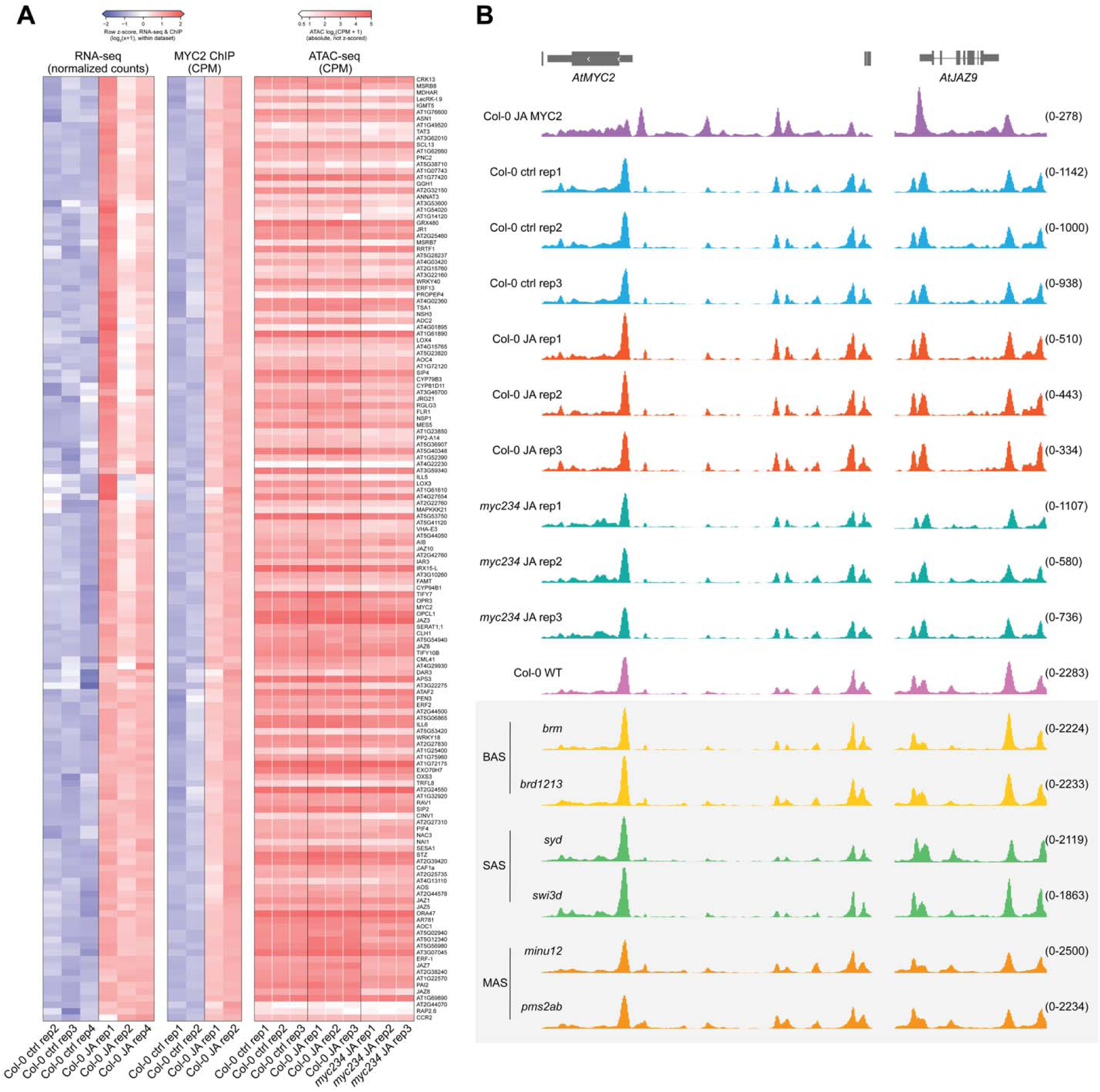
The relationship between the JA accessome, the MYC2 cistrome, and the JA transcriptome A, Heatmap of 145 MYC2 direct target genes, defined as genes nearest to the top 500 MYC2 ChIP-seq peaks (ranked by MACS2 peak score) that are up-regulated after 2 h of JA treatment (log2 FC > 1, adjusted *P* < 0.05). Left, RNA-seq normalized counts in Col-0 seedlings under control conditions and after 2 h of JA treatment (*n* = 3). Middle, MYC2 ChIP-seq signal in control and JA-treated Col-0 seedlings (*n* = 2), shown as reads per million (CPM) within ±100 bp of each MYC2 peak summit; for genes with multiple MYC2 binding sites, signal was averaged across all sites. RNA-seq and ChIP-seq values were log2(x + 1)-transformed and z-scored by row within each dataset, and genes were ordered by hierarchical clustering of the combined z-scores (Euclidean distance, complete linkage). Right, ATAC-seq signal in Col-0 under control and JA conditions and in JA-treated *myc234* seedlings (*n* = 3) at the same MYC2 summit windows, shown as absolute log2 (CPM + 1) values without row normalization, so that accessibility can be compared directly between genotypes, conditions and genes. ATAC-seq data were not used for clustering. **B,** Genome browser screenshots show ACRs at the *MYC2* and *JAZ9* loci. The top track (purple) shows MYC2 ChIP-seq data from JA-treated Col-0 seedlings. All remaining tracks show ATAC-seq: Col-0 under control (blue) and JA (red) conditions, JA-treated *myc234* (teal), and, in the shaded box, data from Guo et al. (2022) (GEO GSE193396) for wildtype (Col-0 WT) and mutants of the three SWI/SNF complex classes: BRM-associated SWI/SNF (BAS; *brm*, *brd1 brd2 brd13*), SYD-associated SWI/SNF (SAS; *syd*, *swi3d*) and MINU-associated SWI/SNF (MAS; *minu1 minu2*, *pms2a pms2b*). All tracks are normalized to sequencing depth.

### Chromatin accessibility at MYC2 targets is regulated through unknown mechanisms

We next asked whether MYCs activity themselves contribute to the pre-existing accessibility of MYC2 target sites. Because MYCs are stabilized by JA, we hypothesized that the low abundance of MYC2 under control conditions, although insufficient for robust detection by ChIP-seq, might nevertheless be sufficient to establish or maintain accessibility at its target sites. To test this possibility, we also performed ATAC-seq in *myc234* mutants following 2 h of JA treatment. Loss of MYC2/3/4 did not result in a global reduction in chromatin accessibility (Fig.3A, B) Moreover, neither the number nor the size of ACRs associated with MYC2 target sites was substantially reduced in *myc234* (Supplementary Table S4). These results indicate that MYC2/3/4 are not required to maintain the accessible chromatin landscape at MYC2 target sites under our tested conditions. Next, we investigated the potential contribution of SWI/SNF chromatin-remodeling complexes to the establishment and maintenance of ACRs at MYC2-binding sites. The *Arabidopsis* BAS, SAS, and MAS chromatin remodeling complexes were previously shown to positively regulate chromatin accessibility, with mutations in individual complex components resulting in the loss of ACRs occupied by several G-box-binding TFs, including PIF4 and ABF1 (Guo et al., 2022). We therefore reanalyzed published ATAC-seq datasets from Col-0 and mutants affecting components of the BAS (*brm* and *brd1/2/3*), SAS (*syd* and *swi3d*), and MAS (*minu1/2* and *pms2a/b*) complexes. MYC2-associated ACRs remained largely unchanged across these mutants indicating that also the BAS, SAS, and MAS complexes are not required to maintain accessibility at these regions (Fig. 3B).

## DISCUSSION

TFs are major drivers of plant stress responses and important targets for agricultural improvement. A fundamental understanding of TF regulomes therefore requires knowledge of their cistromes. However, *in vivo* cistrome-mapping approaches such as ChIP-seq typically require TF-specific antibodies, which remain scarce for plant TFs (Ammari et al., 2026a). Consequently, researchers often rely on transgenic plants expressing epitope-tagged TFs. Although these approaches are widely used and often unavoidable, the resulting cistromes should be interpreted in the context of potentially altered TF dosage (Ammari et al., 2026a). Overexpression constructs can directly elevate TF dosage, but altered dosage can also occur when an epitope-tagged TF is expressed from its native promoter since T-DNA integration occurs at variable genomic locations and can involve multiple transgene copies (Jupe et al., 2019).

Comparison of our wildtype MYC2 cistromes with previously published transgenic MYC2 cistromes revealed that, although targeting of core JA pathway components was broadly conserved, the published datasets derived from epitope-tagged *MYC2* transgenes generally contained more MYC2-binding sites. Differences in experimental conditions, developmental stage, sequencing depth, and analysis pipelines preclude attributing these differences solely to TF dosage. Nevertheless, both the *pMYC2:MYC2-GFP* and *pMYC2:MYC2-YPet* transgenes are expressed in the Col-0 background and therefore contribute epitope-tagged MYC2 in addition to the endogenous MYC2 pool. The *pMYC2:MYC2-FLAG* construct was expressed in a *myc2* mutant background, yet this line also showed substantially greater MYC2 occupancy than wildtype under JA conditions and in most immunoblot experiments, also higher MYC2 protein levels. The most striking difference was however observed under control conditions: whereas endogenous MYC2 displayed only marginal genomic occupancy in Col-0 seedlings, *pMYC2:MYC2-FLAG* seedlings revealed extensive basal MYC2 binding. Our immunoblot analysis further revealed elevated MYC2-FLAG levels already under control conditions, which correlated with the extensive, JA-like cistrome observed in the absence of JA treatment. It’s possible that altered basal MYC2 abundance in *pMYC2:MYC2-FLAG* seedlings could be further amplified by the autoregulatory behavior of MYC2, as MYC2 binds to its own regulatory region and thereby has the potential to reinforce its own expression.

Intriguingly, despite the relatively stable number of peaks, peak heights were substantially smaller in control cistromes of *pMYC2:MYC2-FLAG* seedlings, whereas peak widths remained largely unchanged. We observed a similar trend in Col-0, although to a much smaller extent, with the average occupancy profile showing only a weak indication of a MYC2 peak. Similar peak widths under control and JA conditions are consistent with occupancy of a comparable set of genomic regions. In contrast, differences in peak height may reflect several, non-mutually exclusive scenarios. First, at bulk level, MYC2 binding in control samples may occur in only a subset of cells, whereas JA treatment increases the proportion of cells exhibiting MYC2 occupancy. Second, MYC2 stability may influence its DNA residence time; shorter residence times under control conditions could reduce the probability of efficient crosslinking of MYC2– DNA complexes, leading to lower ChIP signal intensity, thus a smaller peak but with a similar width. In this scenario, the same set of loci is, on average, bound across conditions, but MYC2 occupancy is more transient in the absence of JA.

The latter model is consistent with previous findings that MYC2 stability is regulated by JA (Zhai et al., 2013), raising the possibility that MYC2 is present at its binding sites under control conditions but sufficiently short-lived to limit its capture by ChIP-seq. However, most previous assessments of MYC2 stability were performed using *MYC2* overexpression lines (Zhai et al., 2013, Jung et al., 2015, Chico et al., 2020, Shin et al., 2012, Chico et al., 2014), and the extent to which these observations translate to endogenous MYC2 in wildtype plants remain unclear. We speculate that the JA-induced increase in MYC2 occupancy most likely reflects a combination of increased MYC2 abundance and enhanced MYC2 stability.

Current models generally depict MYC2 as DNA-bound under non-inducing conditions and interacting with the restrictive TPL corepressor complex via JAZs (Zander et al., 2026). Our cistrome data suggest that this may not represent the predominant state of endogenous MYC2 genome-wide. Instead, low MYC2 abundance and rapid turnover in the absence of JA could be sufficient to maintain low steady-state occupancy and prevent activation of MYC2 target genes. Our analysis of TPR1 ChIP-seq under control conditions nevertheless indicates that TPR1 is present at many MYC2 target sites (Ammari et al., 2026b), suggesting that corepressor-associated regulation may operate at these loci even when MYC2 occupancy is difficult to detect. An additional, non-mutually exclusive possibility is that JAZ-bound MYC2 complexes are less efficiently captured by ChIP-seq, which could contribute to the low apparent MYC2 occupancy under control conditions in Col-0 seedlings. Arguing against this possibility, however, is the efficient capture of the MYC2 cistrome under untreated conditions in *pMYC2:MYC2-FLAG* seedlings. Together, these observations suggest that even relatively modest changes in MYC2 abundance can substantially alter the apparent cistrome, particularly under conditions in which endogenous MYC2 abundance and occupancy are normally low.

A major remaining question is how the chromatin landscape contributes to this striking JA-induced expansion of the MYC2 cistrome. MYC2 cistrome expansion is accompanied by extensive transcriptional reprogramming and based on our previous findings, substantial reprogramming of the epigenomic landscape (Choudhary et al., 2024, Ammari et al., 2026b). Both transcriptional and chromatin changes are largely dependent on MYCs; however, how chromatin accessibility contributes to MYC cistrome expansion remains poorly understood. Surprisingly, our ATAC-seq analysis revealed that JA treatment had only a modest effect on the overall accessome under our experimental conditions, with quantitative changes detected at only 1,651 DARs. This relative stability contrasts with the highly dynamic MYC2 cistrome, transcriptome, and histone modification landscape following JA perception, indicating that large-scale changes in chromatin accessibility are not required for MYC2 cistrome expansion.

A largely pre-established accessome may be a more general feature of environmentally responsive transcriptional networks. A similar phenomenon has been observed during the *Arabidopsis* low R:FR light response, where the overall accessible chromatin landscape changes only modestly and most PIF7-binding sites are already accessible before PIF7 recruitment (Paulisic et al., 2025). Related observations have been made in mammalian systems, including the glucocorticoid receptor regulome, where many stimulus-responsive TF-binding sites reside within chromatin that is accessible before activation (John et al., 2011) Importantly, similar ATAC-seq profiles between two conditions do not necessarily imply an unchanged regulatory landscape, as the proteins occupying accessible regions can change substantially without producing major alterations in overall accessibility. Thus, a comparatively static accessome can support highly dynamic TF occupancy and transcriptional responses.

Given the low abundance and stability of MYC2 under control conditions, we initially hypothesized that transient occupancy by MYCs might nevertheless contribute to maintaining accessibility at MYC2 target sites. However, our analysis of JA-treated *myc234* plants argues against this model. MYC2 target regions remained accessible in the absence of MYC2/3/4, with no widespread loss of MYC2-associated ACRs. Other G-box-binding factors, including MYC5, potentially contribute to maintaining accessibility in the absence of MYC2/3/4, although such redundancy is unlikely to fully account for the remarkably stable accessibility landscape. Consistent with this conclusion, our reanalysis of published ATAC-seq datasets from untreated *Arabidopsis* mutants affecting the MAS, SAS, and BAS chromatin-remodeling complexes likewise revealed no widespread loss of accessibility at MYC2-bound ACRs.

Based on these findings, we propose that JA-responsive transcription operates within a largely pre-configured chromatin accessome. This pre-configuration is regulated through an unknown mechanism. Rather than requiring MYC2 to establish ACRs following JA perception, MYC2 exploits a pre-existing permissive accessome. JA-induced increases in MYC2 dosage and stability can therefore drive rapid expansion of the MYC2 cistrome across pre-accessible regulatory regions, followed by extensive transcriptional and epigenomic reprogramming. We propose that this separation between a relatively stable accessome and a highly dynamic cistrome provides a mechanism for maintaining regulatory regions in a poised state, enabling rapid TF recruitment and transcriptional reprogramming in response to environmental signals.

## MATERIALS AND METHODS

### Genetic material and growth conditions

All tested mutants and transgenic material used in this study are in the *Arabidopsis thaliana* Columbia-0 (Col-0) background. *myc2* (*jin1-8, SALK_061267*) (Lorenzo et al., 2004), *myc2 myc3 myc4* (Fernandez-Calvo et al., 2011), and the *myc2 MYC2:MYC2-FLAG* line (Hou et al., 2010) were described previously. Seeds were surface sterilized with bleach, rinsed with water, and stratified in the dark for 2 days at 4°C before being planted on agar plates containing Linsmaier and Skoog (LS) medium (Phytotech Labs), supplemented with 1% sucrose (Sigma), and grown in a growth chamber (AR-41L3; Percival scientific) with a 16 h light/8 h dark cycle, at 19°C. Ten-day-old seedlings were treated with gaseous methyl jasmonate (MeJA; 95% purity; Sigma-Aldrich) in closed transparent containers by applying MeJA to Whatman paper at a concentration of 1 μL per 1 L of container volume. Following MeJA treatment for the indicated times, tissue was rapidly harvested, weighed, and subjected to crosslinking. Tomato (*Solanum lycopersicum* cv. Micro-Tom) seedlings were grown in soil under a 16-h light/8-h dark photoperiod and treated with MeJA using the same setup.

### Cloning of MYC2 constructs

For the generation of *myc234 MYC2:MYC2-4xMYC* plants, the *MYC2* coding sequence including 1949 bp of the regulatory region (fwd. primer, 5′-CACCTTGGTCACATACAAATTTGGATCAGG-3′; rev. primer, 5’-ACCGATTTTTGAAATCAAACTTGCTCTG-3’) was PCR-amplified and purified using the QIAquick PCR & Gel Cleanup Kit (QIAGEN) using *Arabidopsis* Col-0 genomic DNA. The *pMYC2-MYC2* fragment was introduced into pENTR/D-TOPO using directional TOPO cloning (Thermo Fisher Scientific), transformed into *E. coli* DH5α cells, and positive clones were confirmed by restriction digestion and sequencing. Subsequently, the *pMYC2-MYC2* fragment was introduced via Gateway LR reaction using LR clonase II (Thermo Fisher Scientific) into *pGWB516* (Addgene #74858) resulting in *pGWB516-pMYC2-MYC2-4xMYC* which was subsequently transformed into *myc234* mutants using floral dip (Clough and Bent, 1998). To produce recombinant MYC2-4xMYC protein, the *MYC2* coding sequence without the stop codon was first PCR-amplified and purified with the QIAquick PCR & Gel Cleanup Kit (QIAGEN) using *Arabidopsis* Col-0 cDNA. The purified PCR product was cloned into the Gateway pDONR207 entry vector using BP Clonase II (Thermo Fisher Scientific), transformed into *E. coli* DH5α cells, and positive clones were confirmed by restriction digestion and sequencing. The *MYC2* coding sequence was subsequently transferred into *pGWB516* via LR reaction to generate *pGWB516-MYC2-4xMYC*. The *MYC2-4xMyc* fragment was then PCR-amplified, digested with NdeI and NotI, and ligated into the corresponding sites of *pET28a* using T4 DNA ligase. Following transformation into DH5α cells, positive clones were identified by colony PCR and confirmed by DNA sequencing. The resulting *pET28a-MYC2-4xMyc* construct was used for commercial recombinant protein expression by GenScript.

### Western blotting

Plant samples were harvested and flash-frozen in liquid nitrogen. Frozen plant material was ground to a fine powder using a mortar and pestle, and 200 mg of tissue per sample was used for protein extraction. Ground tissue was resuspended in 2× NuPAGE LDS Sample Buffer supplemented with 2.5% β-mercaptoethanol and heated at 70 °C for 10 min. Samples were subsequently centrifuged at 13,000 rpm for 2 min, and the supernatant was used for SDS-PAGE on NuPAGE 4–12% Bis-Tris protein gels (Thermo Fisher Scientific). Immunoblotting was performed using rabbit polyclonal anti-MYC2 (#R3500-3, Abiocode), mouse monoclonal anti-FLAG M2 (#F1804, Sigma-Aldrich), and rabbit polyclonal anti-Histone H3.1 (#SAB4500353, MilliporeSigma) primary antibodies. Goat anti-rabbit IgG (H+L)-HRP (#1706515, Bio-Rad) and goat anti-mouse IgG (H+L)-HRP (#1706516, Bio-Rad) were used as secondary antibodies.

### ChIP-Seq

ChIP-seq was carried out as a combination of standard ChIP-seq and PHILO ChIP-seq modifications (Choudhary et al., 2024, Kaufmann et al., 2010). In brief, 1 g of plant tissue was used for chromatin extraction following a standard ChIP-seq protocol (Kaufmann et al., 2010). Extracted chromatin was fragmented by 30 cycles of sonication (30 s on, 30 s off) using a Bioruptor Plus (Hologic Diagenode) and subsequently processed using the PHILO ChIP-seq pipeline, including library preparation (Choudhary et al., 2024). We used the following antibodies (5 μl per sample) for our study: anti-MYC2 (#R3500-3, Abiocode), anti-FLAG (#F1804, Sigma-Aldrich). IgG (#015-000-003, Jackson ImmunoResearch) served as a control. 10 μl of Dynabeads Protein A (ThermoFisher Scientific) were used per ChIP sample.

### RNA-seq

For the JA time-series RNA-seq experiments, total RNA was extracted from two-week-old untreated and JA-treated soil-grown *Arabidopsis thaliana* Col-0 plants using the RNeasy Plant Mini Kit (Qiagen). Strand-specific cDNA libraries were generated from 1 µg of total RNA using the TruSeq Stranded mRNA Library Prep Kit (Illumina) according to the manufacturer’s instructions. Sequencing was performed by Admera Health using an Illumina NovaSeq X Series 10B flow cell, generating 2 × 150-bp paired-end reads.

### ATAC-seq

ATAC-seq was performed as previously described (Hsieh et al., 2024), with minor modifications. Briefly, approximately 5 g of fresh untreated and JA-treated (2 h) two-week-old soil-grown Col-0 plants, as well as 5 g of JA-treated (2 h) *myc234* plants, were collected and processed in 2.5-g batches for nuclei isolation. 100,000 nuclei per sample were used for the tagmentation reactions with 2.5 µL of Tagmentase (Hologic Diagenode). Transposed DNA was purified using the Qiagen MiniElute PCR Purification Kit and eluted in 13 µL. Of the eluted DNA, 12 µL was amplified using the NEBNext High-Fidelity 2X PCR Master Mix (New England Biolabs) in a 50-µL reaction containing 1 µL of unique dual-index (UDI) primers (Hologic Diagenode, Set 1). Sequencing was performed by Admera Health using an Illumina NovaSeq X Series 10B flow cell, generating 2 × 150-bp paired-end reads.

### ChIP-seq analysis

ChIP-seq sequencing reads were aligned to TAIR10 (*Arabidopsis thaliana*), using Bowtie2 (v2.4.1) (Langmead, 2010, Langmead and Salzberg, 2012). Data quality was evaluated based on FRiP scores, the proportion of uniquely mapped reads, the number of detected peaks, and consistency across biological replicates. For each MYC2 ChIP-seq experiment, three biological replicates were analyzed under control conditions and three following 2 h of JA treatment. The following publicity available datasets were used for MYC2 cistrome analyses: (GEO GSE249738; Choudhary et al., 2024, *myc2 MYC2:MYC2-FLAG*, ctrl rep1/rep2 GSM7962245/GSM7962246; 2h JA rep1/rep2, GSM7962247/GSM7962248), (GEO GSE133408; Zander et al., 2020, Col-0 *MYC2:MYC2-Ypet*, 2h JA rep1/rep2/rep3/rep4 GSM3949131/GSM3949132/GSM3949133/ GSM3949134), (GSA PRJCA001012; Wang et al., 2019, Col-0 *MYC2:MYC2-GFP*, 0.5h JA rep1/rep2 CRR032460/ CRR032461). To identify MYC2 binding peaks and MYC2 peak summit regions from MYC2 ChIP-seq data, MACS2 (v2.2.7.1) was used (Feng et al., 2012). Heatmaps and aggregate profiles, were generated using deepTools (v3.5.2) (Ramirez et al., 2014). For the annotation of MYC2 binding sites, ChIPseeker (v1.28.3) was used (Yu et al., 2015). Gene ontology analysis was conducted with ShinyGO (v0.85.1) (Ge et al., 2020). BEDTools (v2.25.0) and SAMtools (v1.3.1) were used to process ChIP-seq samples (Li et al., 2009, Quinlan and Hall, 2010). Motif analyses were conducted with MEME-ChIP (Machanick and Bailey, 2011). MYC2 binding was visualized using the Integrative Genomics Viewer (IGV) (Robinson et al., 2011). All sequencing was performed by Admera Health using an Illumina NovaSeq X Series 10B flow cell, generating 2 × 150-bp paired-end reads.

### RNA-seq analysis

Raw FASTQ reads were quality-filtered and adapter-trimmed using fastp (v1.3.3) with default parameters and paired-end adapter detection enabled (Chen et al., 2018). A genome index was generated with STAR (v2.7.11b) using the *Arabidopsis thaliana* TAIR10 genome and annotation, with --sjdbOverhang set to the read length minus one (Dobin et al., 2013). Trimmed paired-end reads were aligned with STAR using default alignment parameters, generating coordinate-sorted BAM files and per-gene read counts. Gene-level read counts were also quantified with featureCounts (Subread v2.1.1) using the GTF annotation, counting read pairs overlapping exon features and assigning them to genes according to gene_id (Liao et al., 2014). Library strandedness was determined empirically from the STAR strand-specific per-gene counts and confirmed to be reverse-stranded; featureCounts was therefore run with -s 2.

Differential expression analysis was performed using DESeq2 (v1.50.2) in R (Love et al., 2014). Sample relationships were assessed by principal component analysis and hierarchical clustering of Euclidean sample-to-sample distances, both calculated from variance-stabilized counts (vst, blind = TRUE). In total, we had 15 libraries for the final analysis. Genes with fewer than 10 counts in at least three samples were excluded. Size-factor normalization, dispersion estimation, and Wald tests were performed using the default DESeq2 settings, with ∼ condition as the design formula and control as the reference level. Each JA treatment time point was contrasted against control. Log2 fold-change estimates were shrunk using the adaptive shrinkage estimator implemented in ashr (v2.2.63) (Stephen, 2017). Genes with a Benjamini-Hochberg adjusted *P* < 0.05 and an absolute log2 fold-change > 1 were considered differentially expressed.

The principal component analysis shown in the figure was generated using the DESeq2 plotPCA function on variance-stabilized counts from the 500 most variable genes. For the heatmap, the union of genes upregulated at one or more time points (2,635 genes) was visualized using shrunken log₂ fold changes relative to control, with the color scale capped at ±2, using Morpheus (Broad Institute). Rows were hierarchically clustered using Euclidean distance with complete linkage, and columns were ordered chronologically. Volcano plots were generated using VolcaNoseR (Goedhart and Luijsterburg, 2020). Overlaps among up- and downregulated gene sets across time points were visualized as an UpSet plot showing exclusive intersections, using custom Python (v3.12.3) scripts with pandas (v3.0.2) and matplotlib (v3.10.8) (Hunter, 2007). Normalized counts for individual genes were plotted in R (v4.3.3) using ggplot2 (Wickham, 2016) as mean ± SEM of three biological replicates. Read coverage at selected loci was normalized to sequencing depth and visualized using the IGV web browser (Robinson et al., 2011)

### ATAC-seq analyses

Paired-end ATAC-seq reads were assessed for sequencing quality using FastQC (v0.12.1) (Andrews, 2010). Adapter and quality trimming was performed using Cutadapt v5.2 (Martin, 2011), with the Nextera adapter sequence (CTGTCTCTTATACACATCT) removed from both reads, a minimum Phred quality score of 20, and a minimum read length of 20 bp. Trimmed reads were aligned to the *Arabidopsis thaliana* TAIR10 reference genome using Bowtie2 (v2.4.1) with the --very-sensitive option and a maximum fragment length of 2,000 bp (Langmead and Salzberg, 2012, Langmead, 2010). Alignments were processed using SAMtools (v1.24) (Li et al., 2009). Properly paired reads with a mapping quality of ≥30 were retained, and alignments were restricted to the five nuclear chromosomes, thereby excluding reads mapping to the chloroplast and mitochondrial genomes. PCR duplicates were removed using SAMtools (v1.24) fixmate and markdup -r, and the resulting filtered and deduplicated BAM files were sorted, indexed, and used for downstream analyses (Li et al., 2009). Genome-wide ATAC-seq coverage tracks were generated using deepTools (v3.5.1) bamCoverage with a bin size of 10 bp (Ramirez et al., 2014).

For analysis of Tn5 insertion sites, filtered and deduplicated paired-end alignments were shifted to account for the offset introduced by Tn5 transposase using the --ATACshift option in deepTools alignmentSieve (Ramirez et al., 2014). The 5′ end of each shifted alignment was subsequently extracted as a single-base-pair Tn5 insertion site. Genome-wide insertion-site coverage was calculated and normalized to counts per million (CPM) Tn5 insertion events to generate normalized BigWig tracks.

For differential accessibility analysis, reproducible ACRs were defined from replicate peak calls and regions overlapping annotated repetitive sequences were excluded, resulting in a final consensus set of 38,880 ACRs. Paired-end fragments overlapping each ACR were quantified using featureCounts (v2.1.1) (Liao et al., 2014). Raw fragment counts were analyzed with DESeq2 (Love et al., 2014), with library-size differences accounted for using DESeq2 median-of-ratios normalization. Pairwise differential accessibility analyses were performed between JA-treated and control samples, JA-treated *myc234* and JA-treated wildtype samples, and JA-treated *myc234* and control samples. DARs were defined as ACRs with a Benjamini–Hochberg-adjusted *P* value (FDR) < 0.05. Log2 fold changes were calculated from normalized ACR counts, with positive and negative values representing increased and decreased accessibility, respectively. The following publicly available datasets were used for chromatin accessibility analyses: (GEO GSE193397, ATAC-seq SubSeries GSE193396; Guo et al., 2022, seedlings, wildtype rep1/rep2 GSM5799538/GSM5799539; *brm* rep1/rep2 GSM5799540/GSM5799541; *brd1213* rep1/rep2 GSM5799542/GSM5799543; *syd* rep1/rep2 GSM5799544/GSM5799545; *swi3d* rep1/rep2 GSM5799546/GSM5799547; *minu12* rep1/rep2 GSM5799548/GSM5799549; *pms2ab* rep1/rep2 GSM5799550/GSM5799551).

## ACKNOWLEDGEMENTS

We thank Themios Chionis, Kevin Malone, and Albin Schlab for their excellent support in maintaining our growth chambers and greenhouse facilities.

## CONTRIBUTIONS

L.D., M.A., and M.Z. designed the research. L.D., M.A., J.G., and M.Z. performed ChIP-seq, RNA-seq and ATAC-seq experiments. L.D., M.A., and M.Z. analysed ChIP-seq, RNA-seq, and ATAC-seq data and performed bioinformatics analyses. M.G., K.M., and M.Z. performed immunoblots. M.G. and M.Z. cloned expression construct for MYC2. M.A., L.D. and M.Z. prepared the figures and wrote the manuscript.

## SUPPLEMENTARY DATA

### SUPPLEMENTARY FIGURES

**Supplementary figure S1** Immunoblot validation of the native anti-MYC2 antibody

**Supplementary figure S2** Comparison of wildtype and transgenic MYC22 cistrome

**Supplementary figure S3.** Time-resolved transcriptional response to JA in *Arabidopsis* Col-0 seedlings.

### SUPPLEMENTARY TABLES

**Supplementary Table S1.** MYC2 ChIP-seq peaks across native-antibody and published datasets

**Supplementary Table S2.** Jasmonic acid pathway genes in *Arabidopsis thaliana*

**Supplementary Table S3:** Transcriptional response of *Arabidopsis thaliana* Col-0 in a JA time-series

**Supplementary Table S4:** Overview of identified and differential ACRs

## SUPPLEMENTARY DATA

**Supplementary Figure S1.**
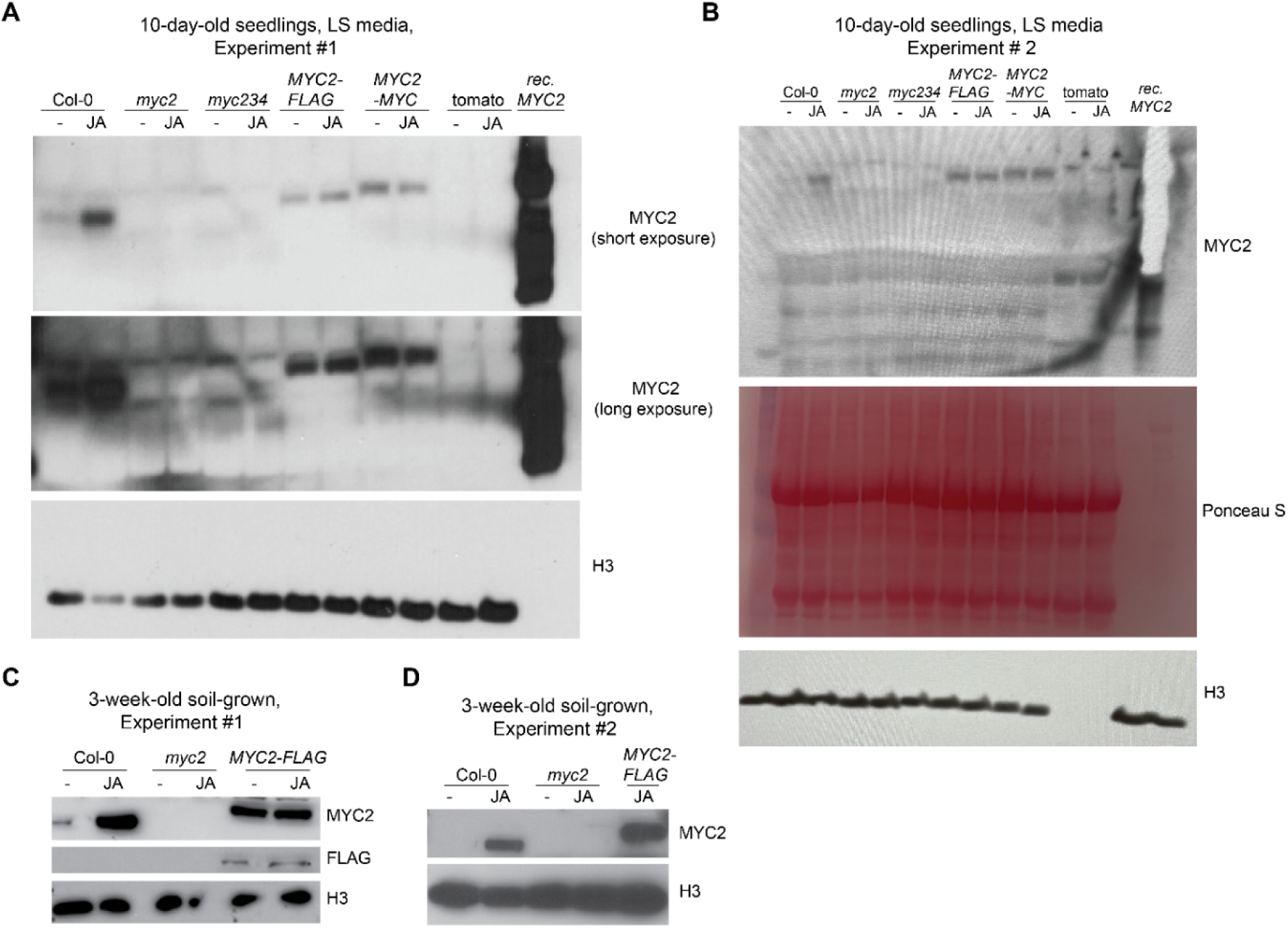
Immunoblot validation of the native anti-MYC2 antibody A, B, Immunoblot analysis of the anti-MYC2 antibody using protein extracts from 10-day-old seedlings grown on LS medium. Two independent experiments are shown. Protein extracts from Col-0, *myc2*, *myc234*, *myc2 pMYC2:MYC2-FLAG*, *myc2 pMYC2:MYC2-4xMYC*, and tomato were analyzed in the absence (−) or presence of JA. Recombinant MYC-tagged MYC2 was included as a positive control but was likely overloaded and yielded a very strong signal. In (**A**), short and long exposures of the MYC2 immunoblot are shown. Histone H3 immunoblotting or Ponceau S staining served as loading controls. **C, D**, Independent immunoblot analyses of protein extracts from 3-week-old soil-grown Col-0, *myc2*, and *myc2 pMYC2:MYC2-FLAG* plants in the absence (−) or presence of JA. MYC2 and FLAG immunoblots are shown as indicated, with histone H3 serving as a loading control.

**Supplementary Figure S2.**
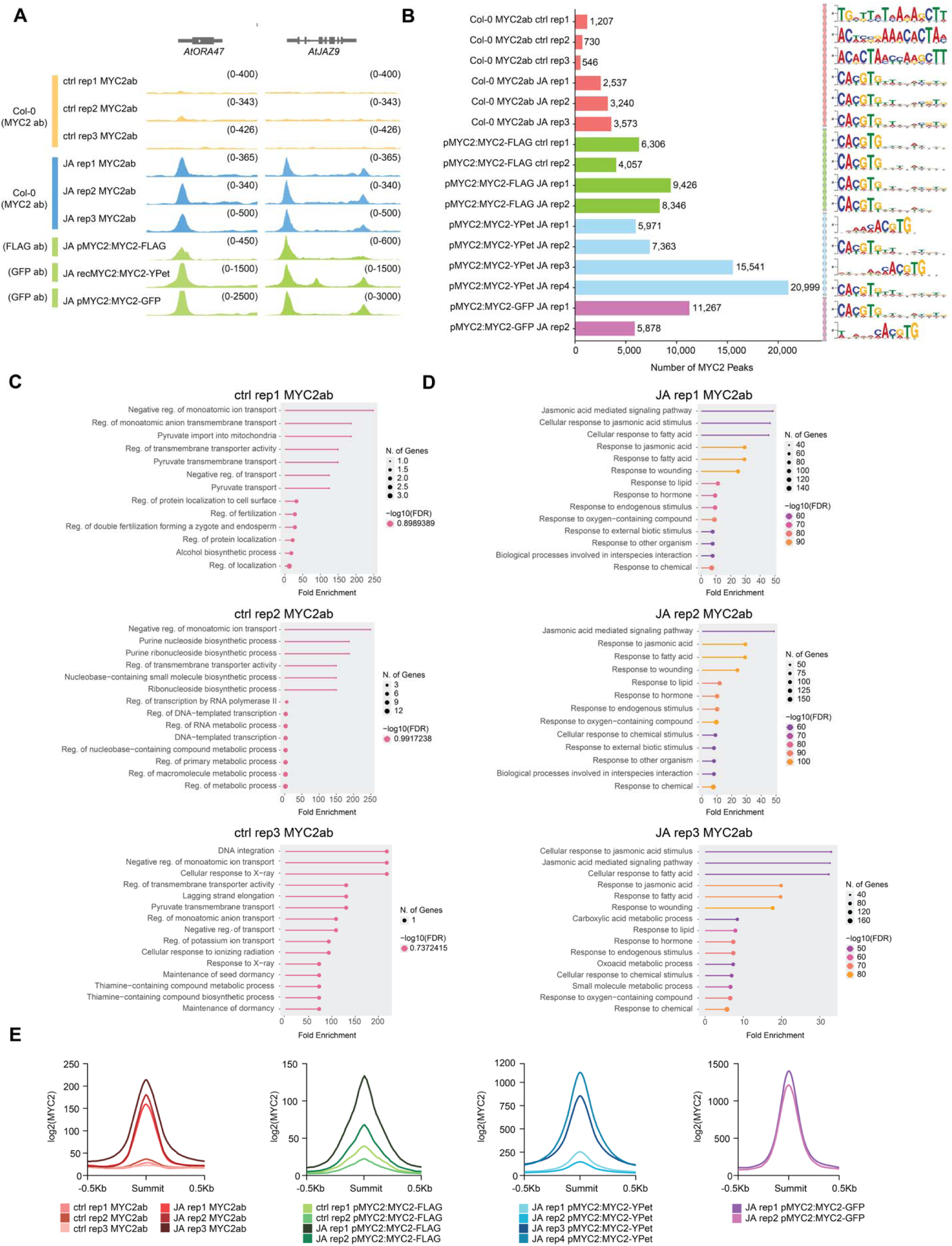
Comparison of wildtype and transgenic MYC2 cistrome A, Genome browser screenshots show MYC2 occupancy at the *ORA47* and *JAZ9* loci in Col-0. MYC2 occupancy in Col-0 under control conditions and 2 h JA treatment was detected with the native MYC2 antibody (blue). MYC2 occupancy from previously published MYC2 ChIP-seq dataset derived from FLAG-, YPet-, and GFP-tagged MYC2 plants are shown in green. **B,** Bar graph shows total number of nuclear MYC2 peaks together with the representative enriched DNA-binding motifs shown for each replicate. Colors indicate the datasets used: native MYC2 antibody (red), *pMYC2:MYC2-FLAG* (green), *pMYC2:MYC2-YPet* (blue), and *pMYC2:MYC2-GFP* (purple). Motif enrichment analysis was performed on the top 500 peak summits selected from MACS2-identified peaks using MEME-ChIP. **C**, **D,** GO enrichment analysis of MYC2 peak-associated genes derived from the top 500 peak summits in Col-0 control replicates (**C**) and JA replicates (**D**). The x-axis represents fold enrichment, point size indicates the number of genes associated with each GO term, and color represents −log10(FDR). **E,** Metagene profiles of MYC2 occupancy across 145 target genes. Colors denote the datasets used: native MYC2 antibody (shades of red), pMYC2:MYC2-FLAG (shades of green), pMYC2:MYC2-YPet (shades of blue), and pMYC2:MYC2-GFP (shades of purple).

**Supplementary Figure S3.**
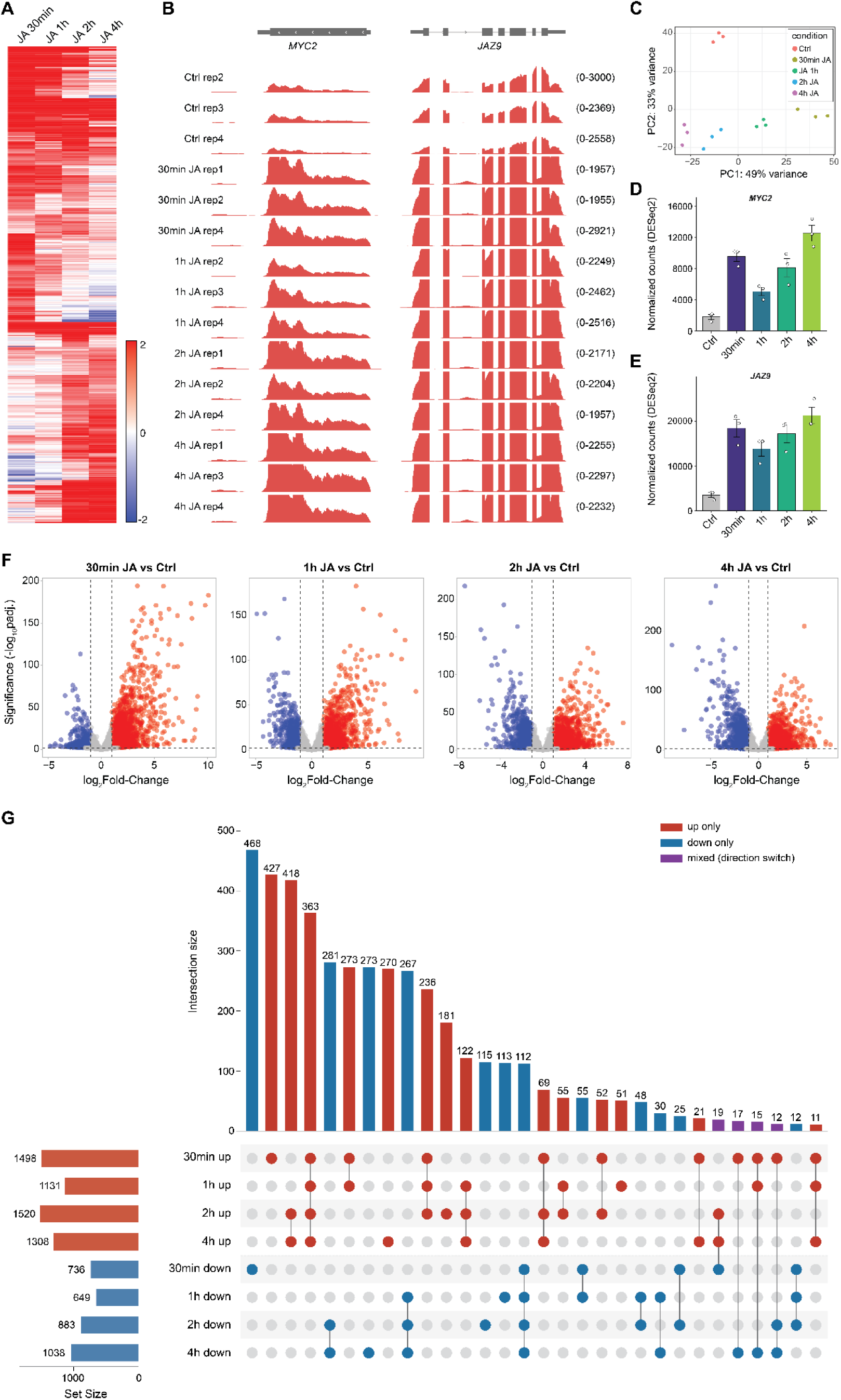
Time-resolved transcriptional response to JA in *Arabidopsis* Col-0 seedlings. A, Hierarchical clustering of the 2,635 genes upregulated (log2 fold-change > 1, adjusted *P* < 0.05) at one or more time points after JA treatment relative to untreated controls. Colors indicate log2 fold-change at 30 min, 1 h, 2 h, and 4 h, with the scale capped at ±2. Rows were clustered using Euclidean distance with complete linkage. **B,** Genome browser screenshots of RNA-seq read coverage at *MYC2* and *JAZ9* for each biological replicate. Coverage was normalized to sequencing depth. **C,** Principal component analysis of variance-stabilized expression data from all 15 libraries (3 biological replicates across 5 conditions). **D, E,** DESeq2-normalized counts for *MYC2* (**D**) and *JAZ9* (**E**). Data are mean ± SEM of *n* = 3 biological replicates, with individual replicates shown as dots. Expression at each time point differed from the control (adjusted *P* < 1 × 10⁻). **F,** Volcano plots showing differential expression at each time point relative to the control. Red, upregulated genes; blue, downregulated genes (log2 fold-change > 1, adjusted *P* < 0.05); gray, not significant. Dashed lines indicate the significance and fold-change thresholds. **G,** UpSet plot of up- and downregulated gene sets across time points. Set sizes are shown on the left. Vertical bars indicate the number of genes in each exclusive intersection, defined by the connected dots. The 30 largest of 41 intersections are shown, encompassing 4,411 of 4,463 differentially expressed genes. Purple bars indicate genes whose direction of change differs between time points.

